# Eco-evolutionary dynamics of temperate phages in periodic environments

**DOI:** 10.1101/2024.07.29.604806

**Authors:** Tapan Goel, Stephen J. Beckett, Joshua S. Weitz

## Abstract

Bacteriophage (viruses that exclusively infect bacteria) exhibit a continuum of infection mechanisms, including lysis and lysogeny in interactions with bacterial hosts. Recent work has demonstrated the near-term advantages of lysogeny over lysis in conditions of low host availability. Hence, temperate phage which can switch between lytic and lysogenic strategies — both stochastically and responsively — are hypothesized to have an evolutionary advantage in a broad range of conditions. To examine generalized drivers of viral strategies over the near- and long-term, we explore the eco-evolutionary dynamics of temperate viruses in periodic environments with varying levels of host availability and viral mortality. We use a nonlinear system of ordinary differential equation to simulate periodically-forced dynamics that separate a ‘within-epoch’ phase and a ‘between-epoch’ phase, in which a (potentially unequal) fraction of virus particles and lysogens survive. Using this ecological model and invasion analysis, we show and quantify how conflicts can arise between strategies in the near-term that may favor lysis and strategies in the long-term that may favor lysogeny. In doing so, we identify a wide range of conditions in which temperate strategies can outperform obligately lytic or lysogenic strategies. Finally, we demonstrate that temperate strategies can mitigate against the potential local extinction of viruses in stochastically fluctuating environments, providing further evidence of the eco-evolutionary benefits of being temperate.

## 1 Introduction

Temperate phages can have two alternative life cycles: lytic and lysogenic. The lytic life cycle involves phage entry into the host, followed by take over of the host machinery to produce new phage and the eventual lysis of the host cell that releases new virions into the environment. These newly released virions can then go on to infect new hosts via horizontal transmission. The lysogenic life cycle involves the integration of the phage genome into the host genome as a prophage, leading to the formation of a lysogen that contains both the phage and the host genome [1–4]. The phage genome reproduces when the lysogenized host cell divides, via vertical transmission. Lysogens can also induce, due to noisy gene regulation and/or cues (extracellular ones such as arbitrium [5–7] or intracellular ones such as stress signals [8]), and re-enter the lytic cycle.

Lysogeny provides direct and indirect benefits to both the phage and the host, in contrast to lysis, where reproductive success of the phage comes at the expense of host mortality. In lysogeny, reproductive success of the phage necessitates reproductive success of the host, since the phage genome replicates with the host genome [9]. More directly, lysogeny can confer superinfection immunity to the infected bacterium [10–12], contribute to horizontal gene transfer and provide novel genetic material to the host [13] and, in cases where the bacterium itself is a pathogen to another organism, increase the pathogenicity of the bacterium [14]. Integrated prophages can also form coalitions with their hosts, allowing for coexistence of multiple phage types with a single bacterial host species [15], potentially contributing to the diversity of phages.

There has been longstanding interest in the evolution and persistence of lysogeny as a strategy for viral reproduction [16]. At first glance, it would seem that lysogeny would have lower reproductive success than lysis: viral replication through lysogeny happens at the rate of cell division and only produces one new viral copy per cell division, while lysis is typically faster and produces several viral copies per lysis event. And yet, temperate phages are abundant in a variety of ecological settings [11]: terrestrial [17], host associated (such as the mammalian gut) [18, 19] and aquatic [20, 21]. In the marine context, lysogen prevalence varies significantly whether estimated based on sequence analysis or prophage induction of isolates from a range of depths, spatial environments and seasons [22, 23]. Metagenomic analysis of dsDNA virus samples isolated from the North Pacific Subtropical Gyre reveals an increase in the prevalence of lysogeny with depth, which correlates negatively with the decrease in host abundance [24]. In polar environments, viral strategies shift from lytic infections during summer periods (when host and resources are abundant) to lysogenic infections during winter periods (when host and resources are relatively depleted) [25].

Theoretical models have been proposed to explain when and why lysogens persist and how their abundance varies with host and nutrient availability. The feast or famine hypothesis suggests that temperate strategies do better when host availability is low and extracellular viral mortality is high [16]. Formalizing the near-term benefits of viral strategies in terms of the basic reproduction number (i.e., the number of newly infected cells produced from a single infected cell in an otherwise susceptible population) have helped reveal how vertical transmission through lysogen replication outperforms lysis when host abundance is low and when extracellular phage mortality is high [26–28]. These models are consistent with observations of a negative correlation between lysogen abundance and host availability in marine environments [24, 25]. Alternative hypotheses, such as Piggyback-the-Winner, suggest the opposite: lysogeny rather than lysis increases with increasing host density [29]. However, counter to this assumption, current implementations of Piggyback-the-Winner models do not explicitly incorporate lysogeny and assume that productivity of viral lysis increases with host density [30] and there is often conflicting (or lack of) evidence to support its claims [30, 31]. The roles and relative value of lytic vs. lysogenic strategies is still a matter of debate (see [29–34]) and may depend on specific ecological contexts. Fluctuations in the availability of hosts, the susceptibility of hosts, and resources available for cells and viruses to complete their life cycle suggest that temperate strategies may function as a means to ‘bet hedge’ against stochastic selection pressures [35]. Indeed, recent conceptual work has emphasized that infections may act along a continuum between lysis and latency [9].

Here, we study the long-term eco-evolutionary dynamics of temperate phages in environments with periodic fluctuations (Figure 1). Notably, periodic changes in host availability and viral decay, generated by external addition of nutrients and hosts, and differential dilution of virions and lysogens, can set conflicting short-term and long-term selection pressures on viral reproductive strategies. These conflicts of selection arise between strategies that do well in the short-term (i.e., within a growth cycle) and those that do well in the long-term (i.e., accounting for survival between growth cycles). Via a combination of pairwise invasion analysis and simulations of stochastically fluctuating environments, we identify a broad range of conditions where temperate strategies outcompete obligately lytic and obligately lysogenic strategies in the long-term, providing further evidence of the eco-evolutionary benefits of being temperate.

**Figure 1.**
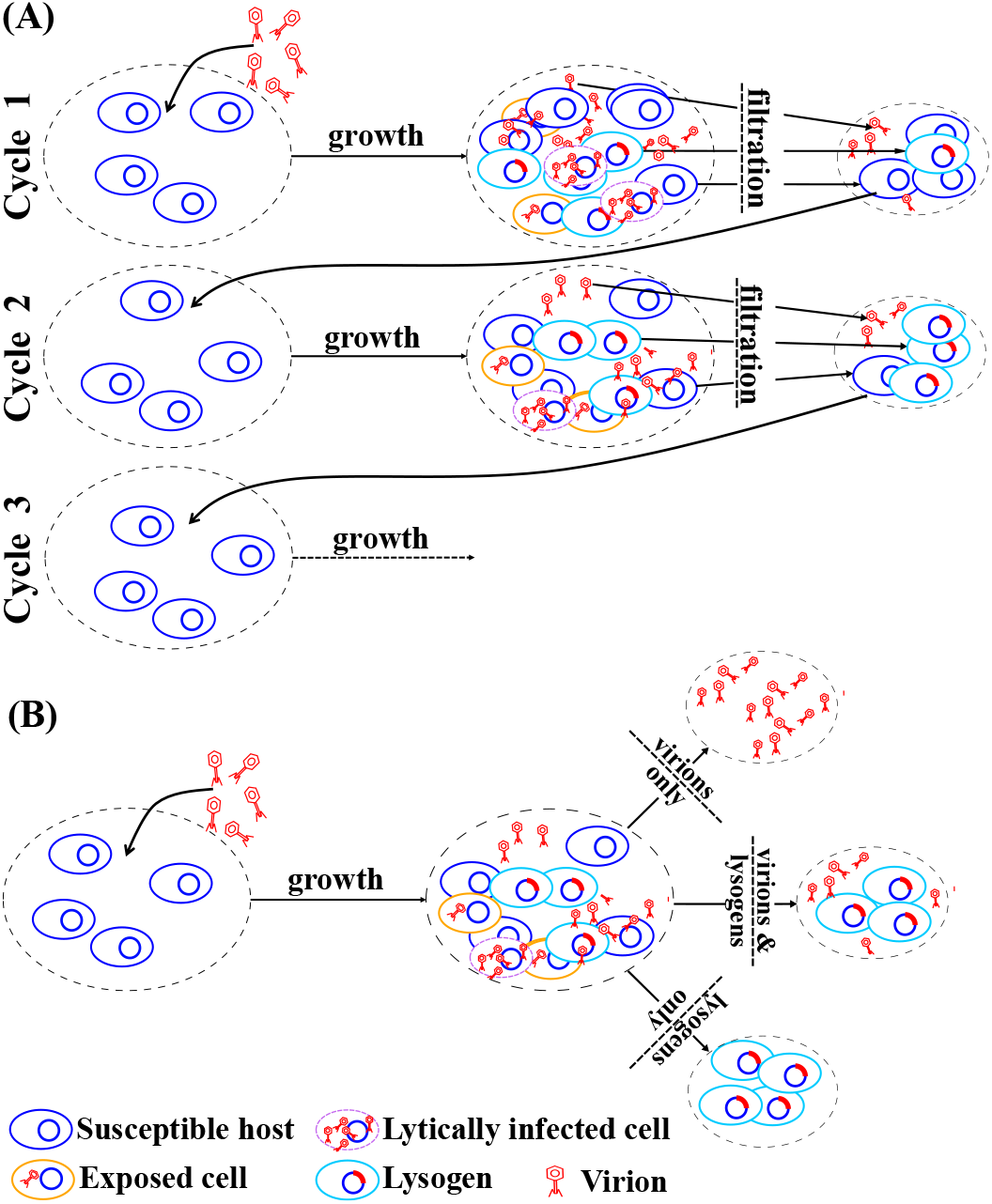
Schematic of the serial passage model: (A) In the first cycle, a fresh batch of nutrients and susceptible cells are inoculated with free virus particles. After a growth cycle, the system will have susceptible hosts, lysogens, lytically infected cells and free virus particles. These cells (viruses) pass through a filter after which only a subset of susceptible cells, lysogens and free viruses remain (in this case half of all susceptible cells, lysogens and free viruses pass through the filter). This filtrate is now used to inoculate the new batch of host cells in cycle 2 and so on. The composition of the system at the end of each growth cycle changes, leading to changes in the composition of the filtrate, which in turn changes the initial conditions for the subsequent cycles. These cycles continue indefinitely or until the system reaches a steady state. (B) The filtration step can isolate virions alone (e.g., via size-dependent filtering), lysogens alone (e.g., via the use of phage-encoded antibiotic selection markers), or a mixture of the two.

## 2 Methods

### 2.1 Resource explicit SEILV model for the population dynamics of a one host-one virus system

We develop an eco-evolutionary model of phage-bacteria dynamics that consists of repeated serial passages. Each passage involves a growth cycle and a subsequent filtration step (see Figure 1). During the growth cycle, phage and bacteria interact in a well-mixed environment. The filtration step occurs at the end of the growth cycle. In the filtration step, different fractions of virions, lysogens, susceptible hosts and resources that remain at the end of the growth cycle are introduced to a fresh batch of resources and susceptible hosts and the next growth cycle begins. This process of coupled growth cycles and filtration steps is then repeated over and over. This serial passage model represents a sequence of boom and bust periods, in which growth occurs during the boom period and viruses and/or cells degrade (potentially at different rates) during the bust periods.

We use a resource explicit SEILV model [26] to describe the phage-bacteria dynamics during a growth cycle. Susceptible hosts (*S*) consume resources (*R*) and replicate, with conversion efficiency *e*. Resource consumption and the consequent increase in host density is assumed to follow Monod kinetics [36] given by the function *ψ*(*R*), where *µ*_max_ is the maximal growth rate of the cells and *R*_in_ is the resource density at which cells grow at half their maximal growth rate. Virions (*V*) adsorb onto cells at adsorption rate *ϕ* and infect susceptible cells. Infected cells are initially classified as exposed cells (*E*). Exposed cells transition at rate *λ* and either form lysogens (*L*) with integration probability *p*, or form lytically infected cells (*I*) with probability (1 − *p*). Lysogens also consume resources and replicate, and they spontaneously induce to a lytic fate at a per capita rate *γ*. The lytically infected cells lyse at a rate *η*, releasing *β* virus particles per lysis event. Susceptible, exposed, lytically infected and lysogenic cells decay at per capita rates *d*_*S*_, *d*_*E*_, *d*_*I*_ and *d*_*L*_ respectively. Virions decay at a per capita rate *m*. These ecological interactions are described by the following set of ordinary differential equations:

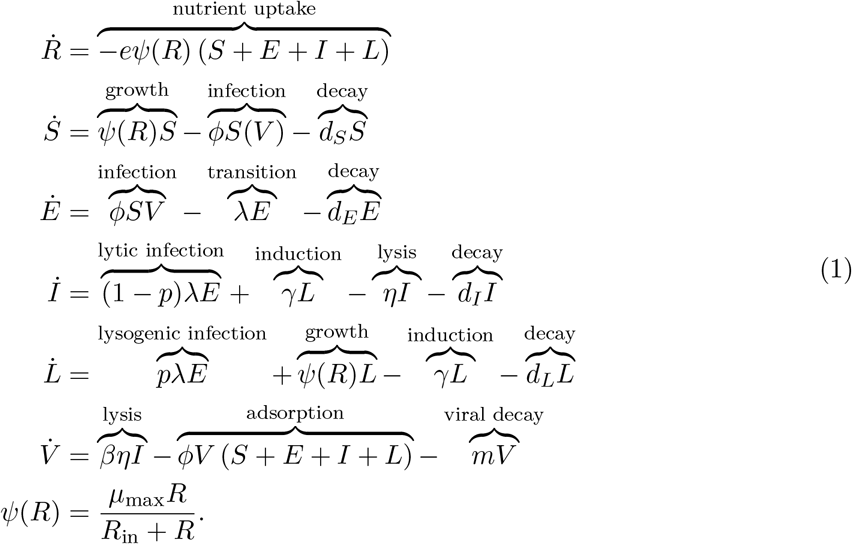

The definitions and values of the parameters in Equation 1 are provided in Table S1 and S2. For convenience, we use matrix notation to describe the state of the system and the filtration steps. The state of the system at time *t* after the beginning of *n*-th growth cycle is given by the 6 dimensional vector 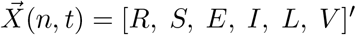, where (*′*) denotes a transpose. Each of the 6 state variables denote densities (here, we use units of mL^−1^). The term 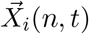 refers to the *i*-th element of the vector 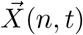.

The filtration step is modelled via matrix multiplication:

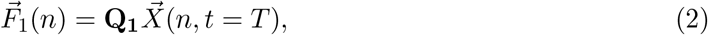

where 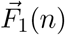 is the filtrate at the end of the *n*-th growth cycle, *T* is the duration of the growth cycle and **Q**_**1**_ = diag(*q*_*R*_, *q*_*S*_, *q*_*E*_, *q*_*I*_, *q*_*L*_, *q*_*V*_) is the filtration matrix. **Q**_**1**_ is a diagonal matrix, with the *q*_*i*_-s being the fractions of each cell type (or virion or resource) that pass through the filter. For example, setting *q*_*V*_ = 0.1 means that 10% of virions left at the end of the *n*-th cycle pass through the filter, setting *q*_*E*_ = 0 means that none of the exposed cells pass through the filter and so on. We set *q*_*R*_ = 0 and *q*_*S*_ = *q*_*E*_ = *q*_*I*_ = 0, and vary *q*_*L*_ and *q*_*V*_ unless noted otherwise. The subsequent growth cycle then starts with the following initial condition:

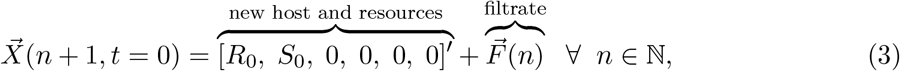

where *R*_0_ = 100 *µ*g/mL and *S*_0_ = 10^7^ cells/mL are fixed densities of resources and susceptible hosts that are added at the beginning of each cycle. We use the following initial conditions for the first growth cycle:

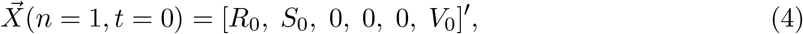

where *V*_0_ = 10^4^ virions/mL is an initial density of virions.

The system of repeated cycles has reached a steady state when:

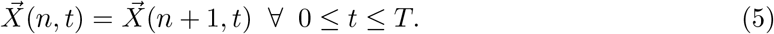

We denote the steady state growth cycle with *n*^*^ and the state of the system at the beginning of the steady state growth cycle with 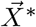:

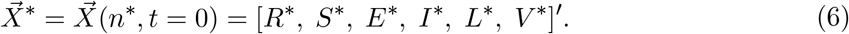

*R*^*^, *S*^*^, *E*^*^, *I*^*^, *L*^*^, *V* ^*^ are the steady state densities at the beginning of the cycle *n*^*^.

### 2.2 Resource explicit SEILV model for the population dynamics of a one host-two virus system for evolutionary invasion analysis

We extend the model for the one host-one virus system to incorporate a second virus. We assume that the two virus types (denoted by *a* and *b*) differ only in the integration probabilities, i.e., probabilities of forming lysogens (*p*-s), and in the spontaneous induction rates (*γ*-s) of their respective lysogens. Each viral type is characterized by a (*p, γ*) pair. We assume the presence of super-infection immunity – virions can adsorb onto infected cells, but they do not take over previously infected cells. The growth cycle dynamics are described by the following ordinary differential equations:

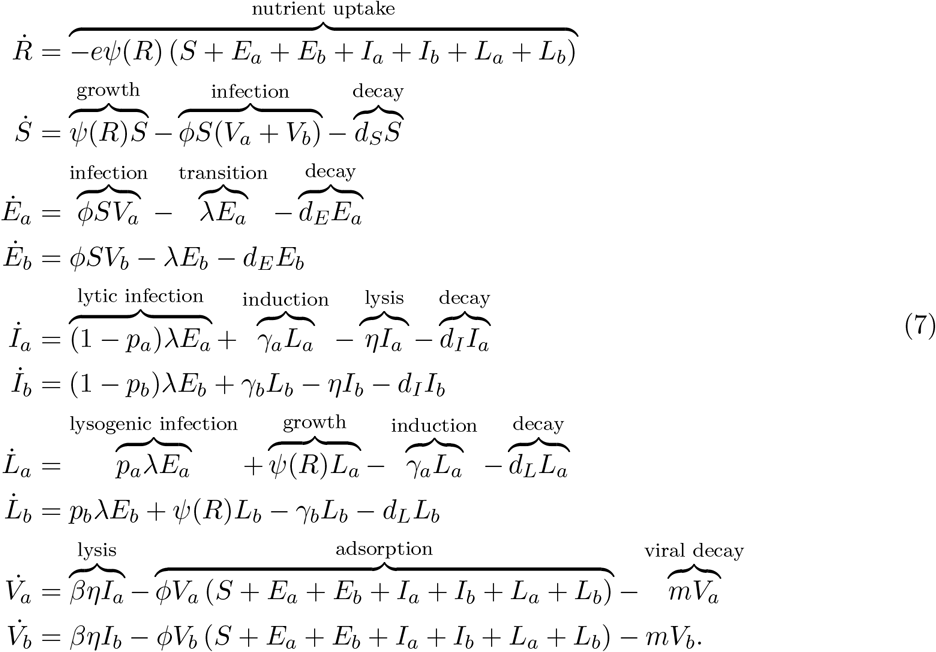

The state of the system at time *t* after the start of the *n*-th growth cycle, is given by a 10 dimensional vector 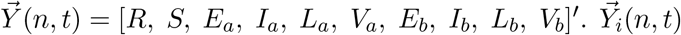 denotes the *i*-th element of the vector 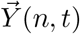. Similar to the model for the one host-one virus system, the filtration step at the end of the *n*-th and initial condition for the (*n* + 1)-th cycle are defined by:

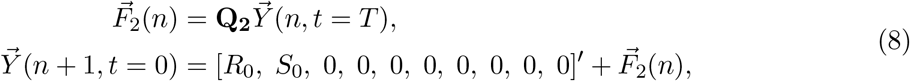

where **Q**_**2**_ = diag(*q*_*R*_, *q*_*S*_, *q*_*E*_, *q*_*I*_, *q*_*L*_, *q*_*V*_, *q*_*E*_, *q*_*I*_, *q*_*L*_, *q*_*V*_) is a filtration matrix similar to **Q**_**1**_.

We use this model to analyze whether a mutant (represented by virus type *b*), appearing at a low initial frequency, with traits (*p*_*b*_, *γ*_*b*_) can invade in a population where the resident (represented by virus type *a*) with traits (*p*_*a*_, *γ*_*a*_) is at steady state with the host. To simulate this scenario, we first let the one host-one virus system with virus type *a* reach steady state 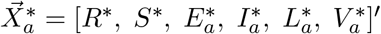. We then initialize the one host-two virus system with the initial condition:

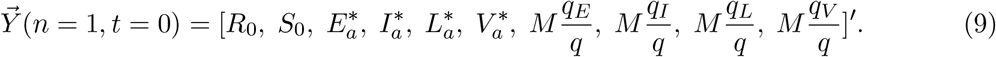

*M* is the (small) initial total density of viral genome copies of the mutant and *q* = *q*_*E*_ + *q*_*I*_ + *q*_*L*_ + *q*_*V*_ is a normalization factor. The normalization ensures that irrespective of the filtration matrix, the total density of mutant viral genome copies introduced into the system is always *M*. Out of the *M* mutant viral genome copies, 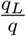 of them are in the form of lysogens, 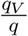 of them are in the form of virions and so on. We say that virus type *b* invades successfully if, in the first few cycles, the total genome density of virus type *b* at the beginning of successive growth cycles increases above *M*. On the other hand, if the total genome density of virus type *b* at the beginning of successive growth cycles decreases below *M*, we say that the invasion failed (see subsection 2.3 for details of the invasion criterion).

### 2.3 Simulation Details

Mathematically, Equation 5 defines the condition for the steady state of the one host-one virus system. Numerically, however, we define the system to have reached a steady state 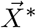 by the *n*^*^-th growth cycle if, in ten consecutive growth cycles starting at *n*^*^, the initial densities of cells (and virions) in successive growth cycle are within a threshold *ϵ* of each other. In practice, we require:

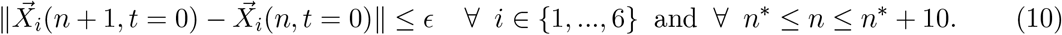

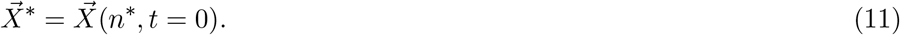

Here, 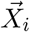 is the *i*-th element of the vector 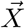 (i.e., 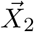 is the density of susceptible cells, 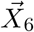 is the density of virions, and so on) and *ϵ* is the critical density threshold. In this study, we assume that the total volume of the system is 1000 mL. Therefore, densities below *ϵ* = 0.001/mL correspond to less than one cell (or virion) in the entire system. So any difference in densities below *ϵ* are not meaningful in practice.

In the invasion analysis we use *M* = 10*ϵ* which corresponds to ten viral genome copies in the entire system, such that the mutant appears at a much lower frequency compared to the resident virus. Further, we only perform invasion analysis when there are more than 100 copies of the resident viral genome, i.e., 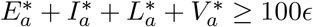 (where 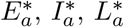 and 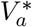 are defined in Equation 9). This is to ensure that the resident is present at a much higher frequency than the mutant and, to ensure that the invasion dynamics are not driven entirely by transients.

To assess if virus type *b* invades successfully, we evaluate whether the total viral genome density of virus type *b* has increased or decreased after *k* growth and filtration cycles. We choose *k* = 10 in practice to reduce the impact of transients. If, after *k* cycles, the total viral genome density of virus type *b* is greater than its initial density *M* and is continuing to increase, then the invasion is classified as successful. On the other hand, if the total viral genome density of virus type *b* is less than *M* and continuing to decrease after *k* cycles, the invasion is classified as unsuccessful. We continue this iterative process until either the condition for invasion success or the condition for invasion failure is met.

Simulation parameters, unless stated otherwise, are provided in Table S1 and Table S2. All code was written in MATLAB 2022b and MATLAB R2023b and, is available for download at https://github.com/tapangoel1994/EcoEvoDynamicsInPeriodicEnvironments and is archived on Zenodo at [37].

## 3 Results

### 3.1 Virus-host dynamics in the near-term

We simulate the SEILV model with one virus and one bacterial host, given conditions for fast-growing bacteria, such as *E. coli* [38]. Figure 2 shows the population dynamics for the first 24 hr growth cycle for three different virus types, each characterized by an induction rate *γ* and integration probability *p*: (a) obligately lytic (*p* = 0, *γ* = 0 hr^−1^), (b) temperate (*p* = 0.5, *γ* = 0.083 hr^−1^) and (c) obligately lysogenic (*p* = 1, *γ* = 0 hr^−1^). The obligately lysogenic virus represents an extreme case where the virus never re-enters its free virus particle life stage. We include it here to evaluate the impacts of distinct life history strategies. Across the three viral strategies considered here, in the first four hours the host density increases since cell division outpaces infection. As the density of virions increases and the density of resources decreases, the susceptible population begins to decline due to the increased rate of infection and the slowing down of cell division (Figure 2). The obligately lytic virus exploits the high density of susceptible hosts and produces new virions through lytic infections, which go on to infect other susceptible hosts. As a result, the virion density increases, slowly at first due to the lack of virions, then quickly due to an abundance of both hosts and virions and slowly again due to the lack of susceptible hosts (Figure 2A). The obligately lysogenic virus, on the other hand, only produces lysogens upon infection, which in turn only reproduce by cell division. First, the lysogen population grows quickly due to lysogenic infections and cell division and then begins to decline due to cell death (Figure 2C). The temperate virus produces both virions and lysogens, albeit at slower rates than the obligately lytic and obligately lysogenic viruses, respectively (Figure 2B). Temperate lysogens produce new lysogens through cell division, and they also produce new virions through induction. These new virions, in turn, can go onto infect additional cells, some of which may become lysogens. As a result, by the end of the growth cycle, the temperate virus can produce more lysogens than the obligately lysogenic virus. So, for a given environmental context, different viral strategies lead to different ecological population dynamics and consequently, different reproductive fitness as measured in terms of the total number of viral genomes produced during a single growth cycle. As shown in Figure 2D, the obligately lytic virus produces the largest number of viral genome copies (the sum of viral genomes inside infected cells and free virions) over the 24 hr growth phase, followed by the temperate and lastly obligately lysogenic viruses. Therefore, when only considering variation in integration and induction over the timescale of a single growth cycle (24 hr) and given abundant available host cells, the obligately lytic virus has greater reproductive fitness than the temperate virus, which in turn has greater reproductive fitness than the obligately lysogenic virus.

**Figure 2.**
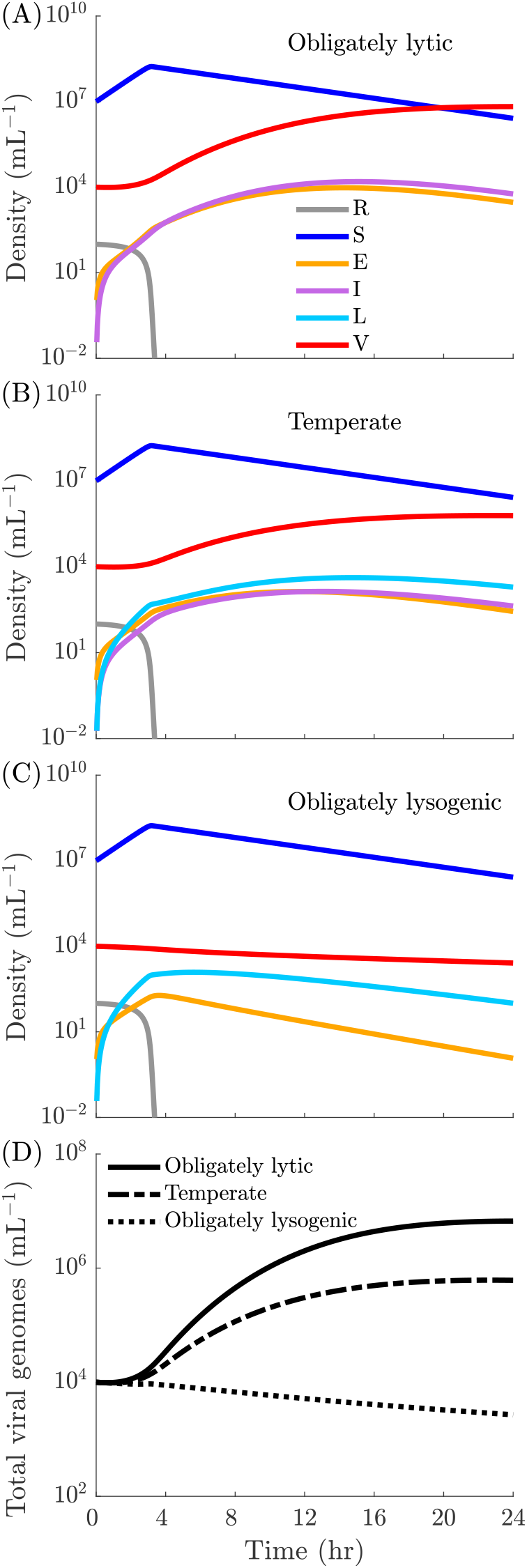
Near-term population dynamics: Population dynamics during a 24 hr growth cycle for (A) obligately lytic (*p* = 0, *γ* = 0 hr^−1^), (B) temperate (*p* = 0.5, *γ* = 0.083 hr^−1^) and (C) lysogenic (*p* = 1, *γ* = 0 hr^−1^) viruses. (D) Total viral genome density (*E* + *I* + *L* + *V*) for the obligately lytic (solid line), temperate (dashed line) and obligately lysogenic (dotted line) viruses. All other simulation parameters are given in Table S1 and Table S2.

### 3.2 Virus-host dynamics in the long-term

We investigate cycle-to-cycle dynamics for different viral strategies in response to different filtration conditions. We consider filtration conditions where (a) only virions, (b) a mixture of virions and lysogens and, (c) only lysogens pass from one cycle to the next. The different filtration conditions favor different viral strategies over the long term: when only virions passage from cycle-to-cycle, we expect strategies that favor lysis to succeed in the long-term – as lysogens are removed at the end of every cycle. On the other hand, when only lysogens pass through the filter, we expect strategies which include lysogeny will succeed – as free viruses will not passage to the next growth cycle. However, it is unclear how cycle-to-cycle filtration interacts with the nonlinear ecological dynamics during the growth cycle in determining the success of viral productive strategies.

First, we consider what happens when only virions passage from one cycle to the next. This scenario simulates the serial passage protocol commonly used in experimental evolution studies of phage virulence. Figure 3A shows the dynamics of total viral genome copies through the first three growth cycles, and Figure 3B shows the longer term cycle-to-cycle dynamics. In the first growth cycle, the obligately lytic virus produces the largest number of viral genome copies, all in the form of virions (Figure S1A). As a result, the obligately lytic virus transfers the maximum number of viral genome copies to the next cycle. The temperate and lysogenic viruses, on the other hand, produce fewer total viral genome copies (Figure S1C, E). Further, only some of these new genome copies are virions (the rest are lysogens) (Figure S1C, E). As a result, the temperate and obligately lysogenic viruses transfer fewer viral genome copies to the next cycle (Figure 3A).

**Figure 3.**
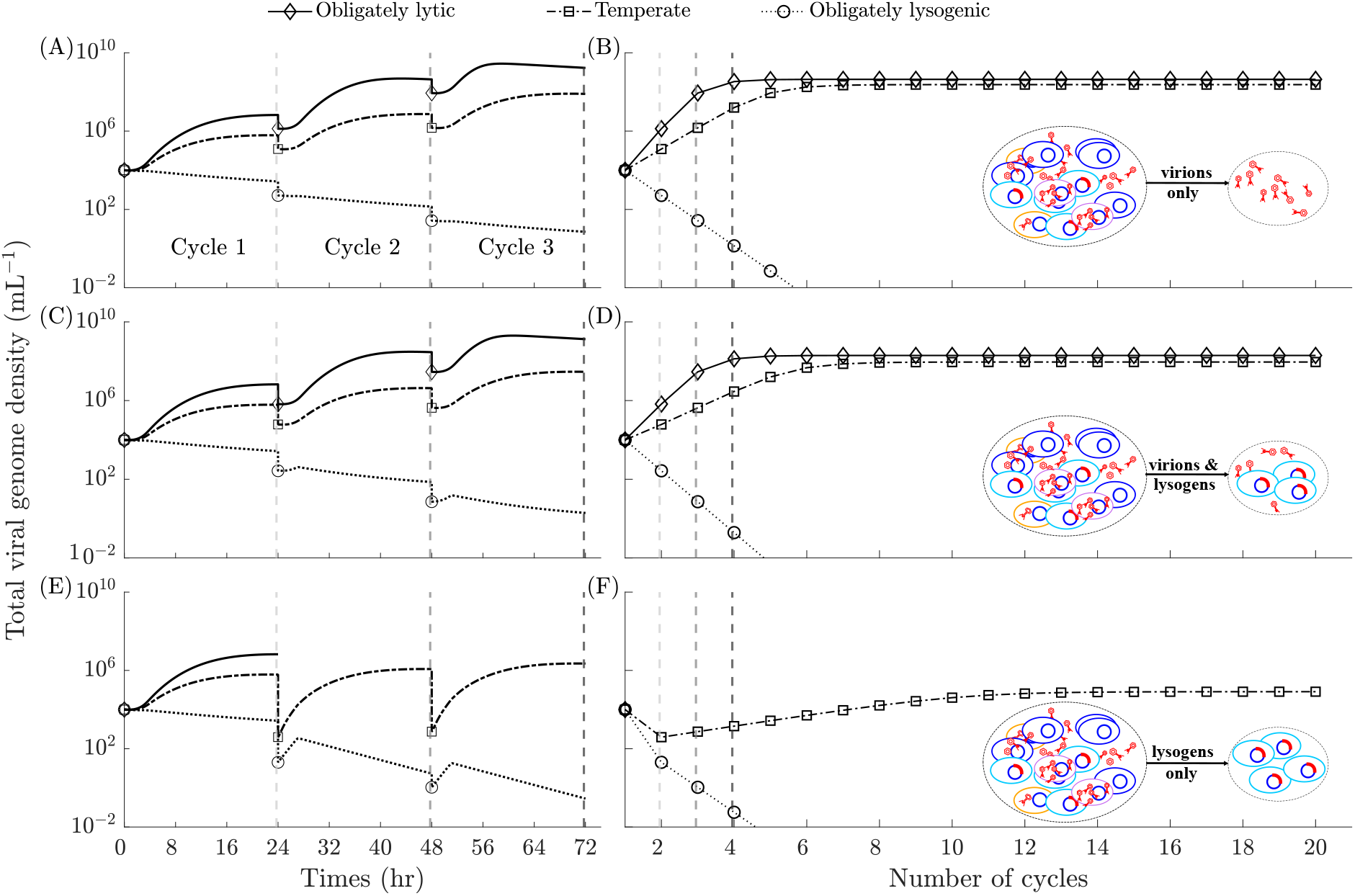
Long-term population dynamics: Total viral genome densities over the first three 24 hr growth cycles for obligately lytic (*p* = 0, *γ* = 0 hr^−1^) (⋄), temperate (*p* = 0.5, *γ* = 0.083 hr^−1^) (□) and lysogenic (*p* = 1, *γ* = 0 hr^−1^) (◦) viruses when (A) only virions (*q*_*V*_ = 0.2), (C) lysogens and virions (*q*_*V*_ = 0.1, *q*_*L*_ = 0.1) and, (E) only lysogens (*q*_*L*_ = 0.2) pass through the filter. Total viral genome densities at the beginning of each growth cycle for obligately lytic (*p* = 0, *γ* = 0 hr^−1^) (⋄), temperate (*p* = 0.5, *γ* = 0.083 hr^−1^) (□) and lysogenic (*p* = 1, *γ* = 0 hr^−1^) (◦) viruses when (B) only virions, (D) lysogens and virions and, (F) only lysogens pass through the filter. The dashed vertical lines at the 24 hr, 48hr and 72hr marks in plots A, C and E correspond to the dashed vertical lines at cycle number 2, 3 and 4 in plots B, D and F respectively.

As this process repeats from cycle to cycle, we observe that the initial density of viral genome copies across growth cycles stops changing over time (Figure 3B). In fact, after 20 growth cycles, the cycle-to-cycle population dynamics for each viral strategy are identical (up to the critical density threshold *ϵ*) (Figure S1), consistent with the system having reached steady state (see subsection 2.3 for details). At steady state, the obligately lytic virus passes on the highest number of viral genome copies from one cycle to the next (Figure 3B). The obligately lysogenic virus on the other hand does not pass any viral genomes to the next cycle: An obligately lysogenic virus does not produce free virions (Figure S1E, F) and therefore the total viral genome density decreases from cycle to cycle (Figure 3A) until there are no viral genomes left (Figure 3B). The temperate virus, however, persists in the population since virions are produced by both, lytic infection and lysogen induction. So, when virions alone are filtered through across growth cycle, the obligately lytic virus and the temperate virus persist over the long-term, while the obligately lysogenic virus rapidly diminishes until it is eliminated.

Next, we consider what happens when only lysogens passage from one cycle to the next (this corresponds to cases where extracellular virions might rapidly decay or degrade outside of growth periods) (Figure 3E, F). As in the previous scenario, we find that the system reaches a steady state after a few growth cycles (Figure 3F). However, in this scenario, the obligately lytic and the obligately lysogenic virus do not persist in the long-term. Even though the obligately lytic virus produces the most viral genomes in the first growth cycle, none of these genome copies make it to the second cycle since only lysogens pass through the filter (Figure 3E, Figure S3A, B). As a result, the obligately lytic virus perishes after the first filtration (Figure S3E, Figure S3A). The obligately lysogenic and temperate viruses, on the other hand, generate lysogens which can pass on the viral genome from one growth cycle to the next and therefore continue to persist past the first growth cycle (Figure S3E, F). However, the number of viral copies passing from cycle to cycle keeps reducing for the obligately lysogenic virus until falling below the critical threshold *ϵ* (Figure 3F); not enough lysogens are generated within the growth cycle to compensate for the filtration. The temperate virus persists given the balance between lysis which produces more virions that infect host cells during the growth cycle with the direct generation of lysogens that passage the viral genome (as a prophage) across growth cycles (Figure S3C, D).

We also consider what happens when both lysogens and virions passage from cycle to cycle (Figure 3C, D). In this scenario, the obligately lytic virus persists due to the production and passaging of virions, while the temperate virus persists due to the production and passaging of virions and lysogens. The obligately lysogenic virus still fails to persist because it is unable to generate sufficient viral copies during the growth cycle to compensate for the filtration at the end of the cycle.

Comparing the long-term population outcomes for the same viral strategy across different filtration conditions, we find that the total initial viral genome density at steady state for the obligately lytic virus is lower when both lysogens and virions pass through (Figure 3D), compared to when only virions pass between growth cycles (Figure 3B). On the other hand, the total viral genome density at steady state for the temperate virus when both lysogens and virions pass through the filter (Figure 3D) is higher than when only virions pass through (Figure 3B). These results demonstrate that the long-term success of viral strategies depends on the interplay between viral reproductive strategy, ecological feedback during growth cycles (i.e., boom periods), and the relative rates of mortality for the different stages in the viral life cycle during filtration (i.e., bust periods).

### 3.3 Temperate strategies maximize viral fitness under conflicting short-term and long-term selection pressures

As we showed in the previous section, depending on the filtration conditions, the success of viral strategies may be discordant between the long term vs. the short term. Analyzing the fitness of different viral life history strategies requires comparing population dynamics over the long-term. We do so by comparing the total viral genome densities at the beginning of steady state growth cycles across the continuum of viral strategies for different filtration conditions. We utilize invasion analysis and pairwise invasibility plots [39] to assess the relative long-term fitness of viral strategies. First, we allow a one host-one virus (the resident virus) system to reach steady state, then add a second virus type (the mutant) with a different life history strategy at very low density and evaluate whether the population of the second virus increases from cycle-to-cycle, over the course of the first few growth cycles (see subsection 2.3 for details).

We first consider what happens when only virions are passaged from one growth cycle to the next. The obligately lytic strategies (*p* = 0) produce the maximum number of viral genome copies at steady state (Figure 4A). Viruses with more temperate strategies (i.e., *p >* 0) also persist at steady state but do not produce as many viral genome copies (Figure 4A). To analyze invasions, we fix the induction rate of the resident and the mutant to be *γ* = 0.083hr^−1^ (we also use the same value of *γ* for invasion analysis in Figure 4D) but allow them to have different integration probabilities. Thus, the life history strategies of the resident-mutant pairs lie along the horizontal red line in Figure 4A. From the PIP, we see that the temperate strategies with more lytic traits (i.e., lower integration probabilities) always invade residents with more lysogenic traits (i.e., higher integration probabilities), but the opposite does not hold (Figure 4B). Thus, we would expect the virus to evolve towards lower and lower integration probabilities, eventually reaching *p* → 0, in agreement with the one-host one-virus analysis in Figure 4A. In this filtration scenario, lysis is favored in the short-term since a high density of susceptible hosts is added in each growth cycle, and lysis is also favored in the long-term since virions (largely produced by lysis) transfer the viral genome between cycles.

**Figure 4.**
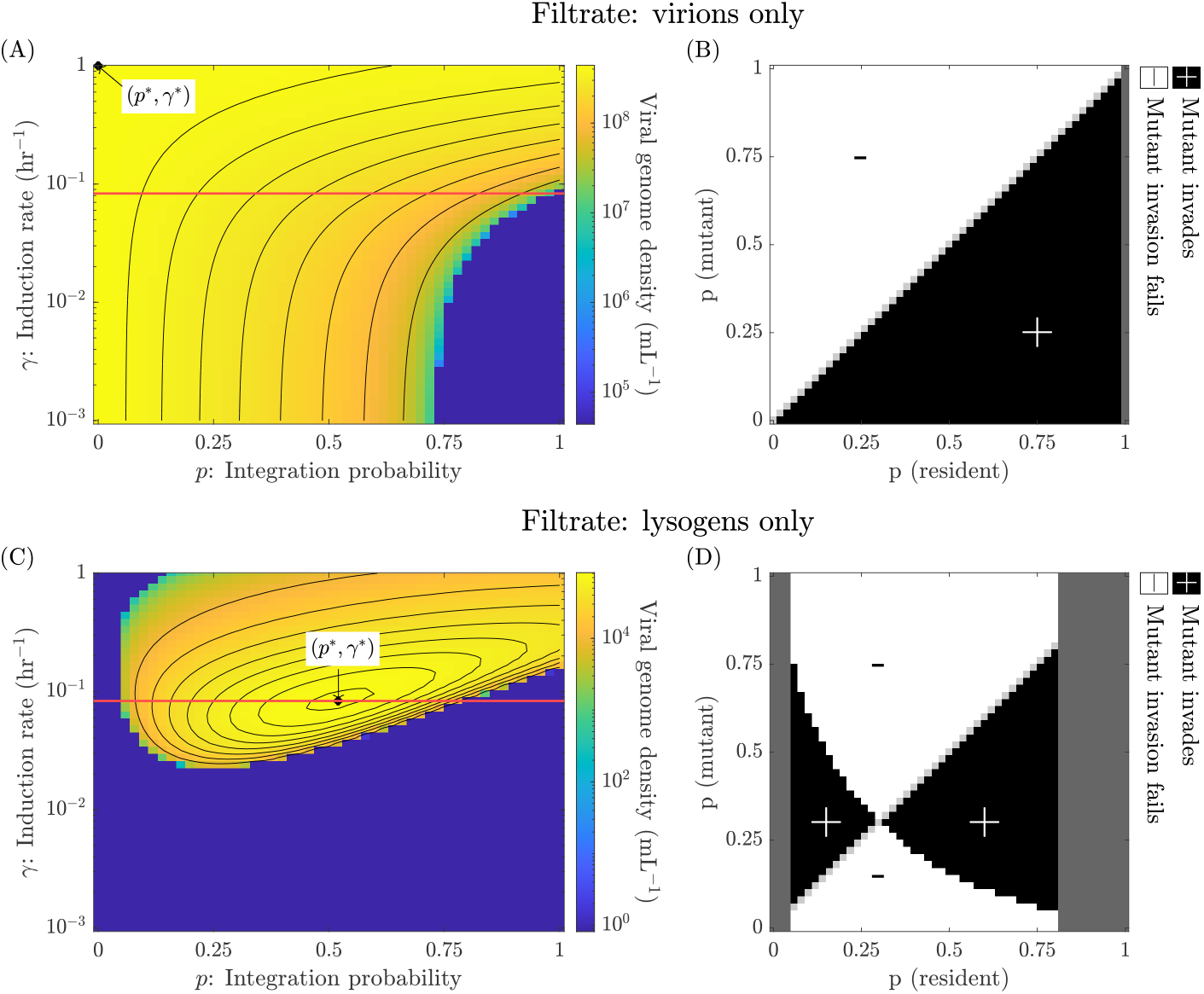
Steady states and invasion analysis for different long-term selection pressures: (A) Heatmap of the density of viral genome copies at the beginning of a steady state growth cycle across different induction rates and integration probabilities when only virions are passaged (*q*_*V*_ = 0.2). (B) Pairwise invasibility plot for viral strategies with different prophage integration probabilities *p* along the horizontal red line (*γ* = 0.832 hr^−1^) in panel (A). Black regions denote successful invasions and white regions denote failed invasions. Dark gray regions correspond to integration probabilities at which the resident virus does not persist in the one host-one virus system (see subsection 2.3 and Figure S5 for details). Light gray marks regions where the resident and mutant have identical life history strategies. (C) Density of viral genome copies at the beginning of a steady state growth cycle when only lysogens are passaged (*q*_*L*_ = 0.2) and (D) the corresponding pairwise invasibility plot. (*p*^*^ = 0, *γ*^*^ = 1 hr^−1^) in panel (A) and (*p*^*^ = 0.52, *γ*^*^ = 0.083 hr^−1^) in panel (C) denote the strategies that maximize the density of viral genome copies being passaged from cycle to cycle at steady state. All other simulation parameters can be found in Table S1 and Table S2.

Next, we consider what happens when lysogens alone are passaged from one growth cycle to the next. Here, the obligately lytic and obligately lysogenic viruses reach extinction while temperate viruses persist (Figure 4C). Within the continuum of temperate viral strategies, some strategies do better than others: we find that an intermediate strategy (*p*^*^ = 0.52, *γ*^*^ = 0.083 hr^−1^) maximizes the steady state viral genome density (Figure 4C). We next consider pairwise invasions for different viral strategies along the *γ* = 0.083 hr^−1^ line (horizontal red line in Figure 4C). The regions *p*_(resident)_ *<* 0.06 and *p*_(resident)_ *>* 0.80 in Figure 4D are infeasible since they correspond to trait values for which the steady state viral genome density of the resident is very small (Figure S5) and therefore, the notion of invasion by a mutant does not arise (see subsection 2.3 for details). In the feasible region 0.06 ≤ *p*(resident) ≤ 0.80, we find an evolutionarily stable strategy (ESS) given by *p*_ESS_ = 0.3 (Figure 4D) that cannot be invaded by a viral mutant using an alternative life history strategy.

This *p*_ESS_ strategy is also convergence stable. Therefore, if the evolutionary timescale is larger than the ecological timescale and mutations are both rare and lead to small changes in the integration phenotype, one would expect a single temperate phage in the system with an integration probability *p* = *p*_ESS_. Similar results have been demonstrated in the context of a host limited model of temperate phages [40].

The success of a temperate strategy in the scenario where lysogens alone are passaged can be explained qualitatively by looking at the conflict between the short- and long-term selection pressures set by the host input and the filtration. Since lysogens alone carry the viral genome from cycle to cycle, the strategy that maintains the highest lysogen density at the end of a growth cycle will maximize the viral genome density at steady state. There are primarily two ways in which new lysogens are generated: lysogens reproducing to make new lysogens (*L* → *L*) and free virions (produced by lysis) infecting susceptible cells, which then form lysogens (*I* → *V* → *E* → *L*) (see [28] for related analysis of temperate life cycles). The first path is only available when resources are present in the system, while the second path is only available when there are susceptible cells in the system. The production of lysogens along the second path depends on having a large number of lytically infected cells, which requires large *γ* for the *L* → *I* transition and low *p* for the *E* → *I* transition. But, maximizing the number of lysogens produced also requires low *γ* to prevent the *L* → *I* transition and high *p* for the *E* → *L* transition. An intermediate temperate strategy that balances this trade-off generates the highest density of lysogens (and therefore viral genome copies) at steady state.

The conflict between the short- and long-term selection pressures leads to another interesting result – when only lysogens passage through the filter, the evolutionary stable strategy (ESS) given by (*p*_ESS_ = 0.3) and the strategy that maximizes the steady state viral genome density (*p*^*^ = 0.52) are different. Notably, the ESS has more lytic traits than the strategy that maximizes the steady state viral genome density. This difference arises from selection operating at two different timescales. While the virus with integration probability *p*^*^ would produce more lysogens in isolation than the virus with integration probability *p*_*ESS*_, when the two viruses compete, the virus with the evolutionarily stable viral strategy (*p*_*ESS*_) infects more hosts in the initial part of a growth cycle as it produces more lytic infections during each cycle. Consequently, there are fewer susceptible hosts available to the strategy with integration probability *p*^*^. Therefore, the ESS does better in the short term and eventually outcompetes the strategy that maximizes the density of viral genomes passaged from cycle to cycle at steady state (Figure S4).

### 3.4 Changing environmental conditions change the eco-evolutionary outcomes of temperate phage-host interactions

To analyze the effect of the growth cycle duration on the success of viral strategies, we focus on the scenario where a fixed fraction of lysogens alone passage from one cycle to the next. We investigate how changing the growth cycle duration changes the density of viral genome copies at the beginning of a steady state growth cycle, as a function of induction rate and integration probability (Figure 5A, C, E). As the growth cycle duration (*T*) increases from *T* = 12 hr to *T* = 24 hr, the space of (*p, γ*) values that allows for viral persistence reduces, as indicated by the increasing purple region in (Figure 5A, C, E). While the induction rate of the strategy that maximizes the viral genome density at the beginning of a steady state growth cycle does not change appreciably as the growth cycle duration increases (*γ*^*^ = 0.095 hr^−1^, 0.083hr^−1^ and 0.083 hr^−1^ for T = 12 hr, 16 hr and 24 hr respectively), the integration probability decreases (*p*^*^ = 1.00, 0.82 and 0.52 for T = 12 hr, 16 hr and 24 hr respectively). To analyze invasions in the context of each of these growth cycle durations, we consider resident-mutant pairs with different integration probabilities but the same induction rate (given by *γ*^*^ = 0.095hr^−1^, 0.083 hr^−1^ and 0.083 hr^−1^ for T = 12 hr, 16 hr and 24 hr respectively). Thus, the resident-mutant pairs lie along the horizontal red lines in Figure 5A, C and E for T = 12 hr, 16 hr and 24 hr respectively. The traits associated with the evolutionarily stable viral strategy become more lytic as the growth cycle duration increases (Figure 5B, D, F): *p*_ESS_ = 0.64, 0.44 and 0.30 for the 12 hr, 16 hr and 24 hr cycle periods respectively. As noted earlier in subsection 3.3, for a given growth cycle duration and filtration condition, we still observe that the evolutionarily stable strategy has a lower integration probability than the strategy that maximizes the viral genome density at the beginning of a steady state growth cycle.

**Figure 5.**
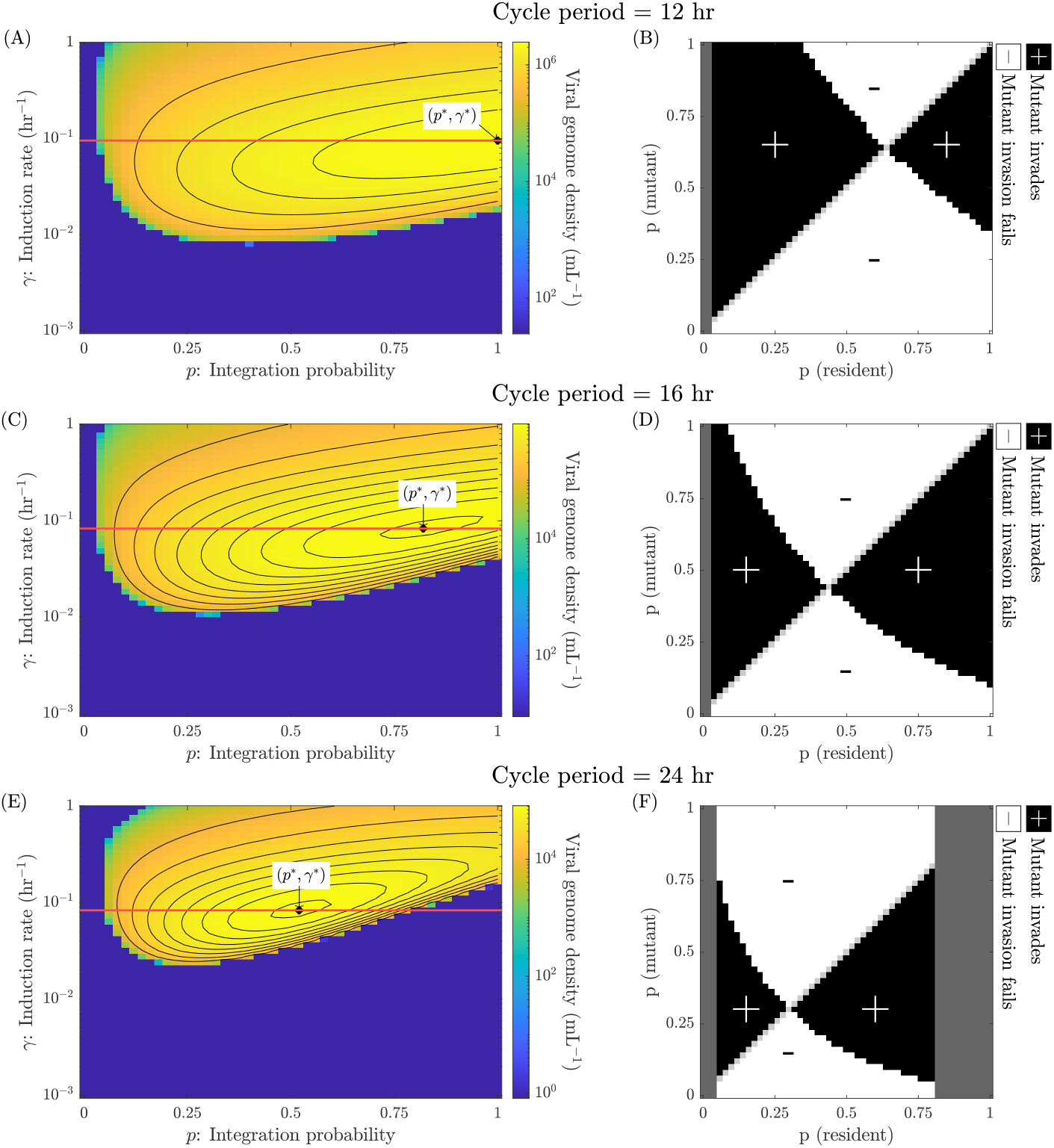
Steady states and invasion analysis for different growth cycle durations: Heatmaps displaying the viral genome density at the beginning of a steady state growth cycle as a function of induction rate and integration probability for (A) 12 hr, (C) 16 hr and, (E) 24 hr growth cycles when only lysogens are passaged (*q*_*L*_ = 0.2). (*p*^*^ = 1, *γ*^*^ = 0.095 hr^−1^), (*p*^*^ = 0.82, *γ*^*^ = 0.083 hr^−1^) and (*p*^*^ = 0.52, *γ*^*^ = 0.083 hr^−1^) mark the strategies that maximize the viral genome density at steady state for the 12 hr, 16 hr and 24 hr growth cycle durations respectively. Panels (B), (D) and (F) are the pairwise invasibility plots for strategies with different integration probabilities *p* for a fixed induction rate given by the horizontal red lines in (A), (C) and (E) respectively. Dark gray regions correspond to integration probabilities at which the resident virus does not persist in the one host-one virus system (see subsection 2.3 and Figure S6 for details). Light gray marks regions where the resident and mutant have identical life history strategies. All other simulation parameters can be found in Table S1 and Table S2.

The link between evolutionarily stable viral strategies and the growth cycle duration can be explained by changes in the relative contributions of the vertical (*L* → *L*) and the horizontal (*I* → *V* → *E* → *L*) transmission modes in production lysogens over time. Initially, when resources are present in the system, the vertical mode of transmission produces lysogens. Once resources are depleted and cell division drops to effectively zero, new lysogens are only produced through infection of susceptible cells via horizontal transmission. Meanwhile, existing lysogens are depleted due to cell death and induction. Since the resources are depleted early on during the growth cycle, the potential for lysogen production via strictly vertical transmission is limited. As growth cycle duration increases, the production of lysogens relies on horizontal transmission for a larger fraction of the growth cycle. Therefore, as the growth cycle duration increases, strategies with a lower integration probability produce more offspring since they favor horizontal transmission. This argument also explains why an increase in the duration of the growth cycle leads to more lytic strategies being favored when only lysogens passage from one cycle to the next, as seen in Figure 5.

However, changing the filtration condition such that only virions passage from cycle to cycle or, some combination of virions and lysogens passage from cycle to cycle, leads to different ecological and evolutionary outcomes. For the two qualitatively different cases where (a) only lysogens pass from cycle to cycle and, (b) only virions pass from cycle to cycle, we obtain qualitatively different relationships between the steady state populations (Figure 4A, C) and the evolutionary outcomes (Figure 4B, D). More quantitatively, given the scenario of passaging lysogens alone, decreasing *q*_*L*_ leads to a smaller region of persistence in the (*p, γ*) parameter space (Figure 6A, C). The strategy that maximizes the viral genome density at steady state when *q*_*L*_ = 0.1 has more lytic traits (*p*^*^ = 0.42, γ^*^ = 0.144 hr^−1^) than the strategy that maximizes the viral genome density at steady state when *q*_*L*_ = 0.2 (*p*^*^ = 0.52, *γ*^*^ = 0.083 hr^−1^). Analyzing invasions between resident-mutant pairs with *γ* = 0.144 hr^−1^ when *q*_*L*_ = 0.1 and, *γ* = 0.083 hr^−1^ when *q*_*L*_ = 0.2, we find that the ESS integration probability remains the same (*p*_ESS_ = 0.30) (Figure 6B, D).

**Figure 6.**
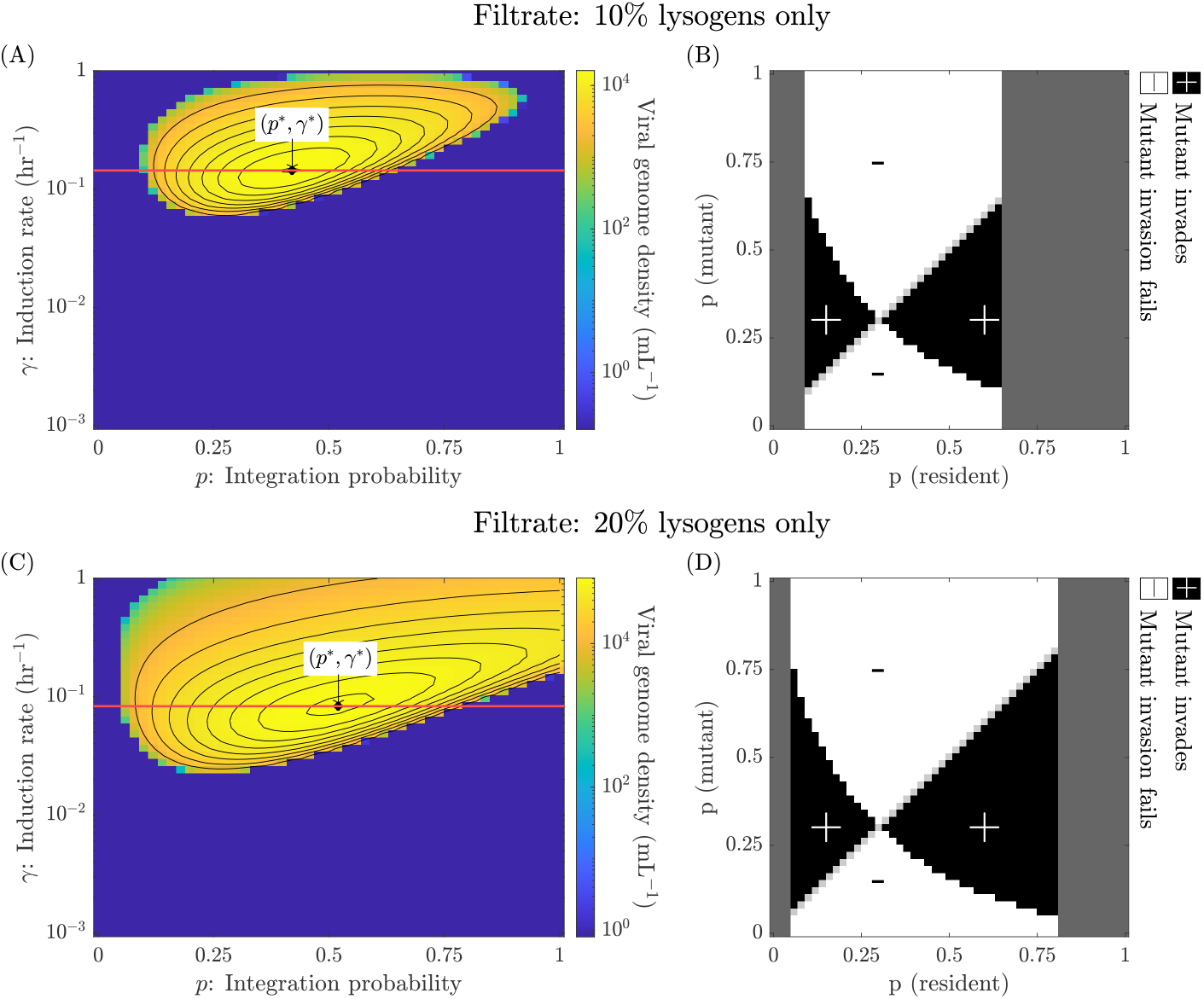
Steady states and invasion analysis for different filtration conditions: Heatmaps displaying the viral genome density at the beginning of a steady state growth cycle as a function of integration probability and induction rate when (A) 10% (*q*_*L*_ = 0.1) and, (C) 20% (*q*_*L*_ = 0.2) lysogens are passaged from cycle to cycle. The duration of the growth cycle in both cases is set to 24 hr. (*p*^*^ = 0.42, *γ*^*^ = 0.144 hr^−1^) and (*p*^*^ = 0.52, *γ*^*^ = 0.083 hr^−1^) mark the strategies that maximize the viral genome density at the beginning of the steady state growth cycle in scenarios corresponding to (A) and (C) respectively. (B) and (D) are the pairwise invasibility plots for viral strategies with different integration probabilities *p* with fixed induction rates, corresponding to the horizontal red lines in panels (A) and (C) respectively. Dark gray regions correspond to integration probabilities at which the resident virus does not persist in the one host-one virus system (see subsection 2.3 and Figure S7 for details). Light gray marks regions where the resident and mutant have identical life history strategies. All other simulation parameters can be found in Table S1 and Table S2.

### 3.5 Being temperate provides protection against environmental stochasticity

Thus far, we have established how temperate strategies evolve and allow for viral persistence in response to short-term and long-term selection pressures in deterministically periodic environments. However, viruses also encounter environmental stochasticity — in the present context this could be represented by changing the duration of growth cycles, the filtration conditions, or both. To explore the impact of environmental stochasticity on long-term viral strategies, we chose to simulate a hundred growth cycles of 24 hr each with 10% lysogens passaging from one cycle to the next while randomly selecting a fraction *q*_*V*_ of virions to be passaged from a log uniform distribution with a span [10^−5^, 10^−1^]. We find that both obligately lytic and obligately lysogenic strategies lead to viral extinction after a few cycles (Figure 7). The obligately lysogenic virus dies out because it does not produce enough viral copies to sustain the virus population across cycles. The obligately lytic virus, on the other hand, dies out more slowly. Local extinction of the obligately lytic virus occurs when stochastic fluctuations generate a series of filtration steps with very low values of *q*_*V*_. The temperate virus, however, overcomes both these problems — through horizontal transmission it produces enough viral copies to prevent gradual extinction and through vertical transmission, it maintains enough lysogens that serial passage conditions representing high virion decay (i.e., low *q*_*V*_ values) do not impact the viral population significantly. This finding further reinforces that being temperate provides robust evolutionary benefits in both deterministic (Figure 3) and stochastically (Figure 7) fluctuating environments.

**Figure 7.**
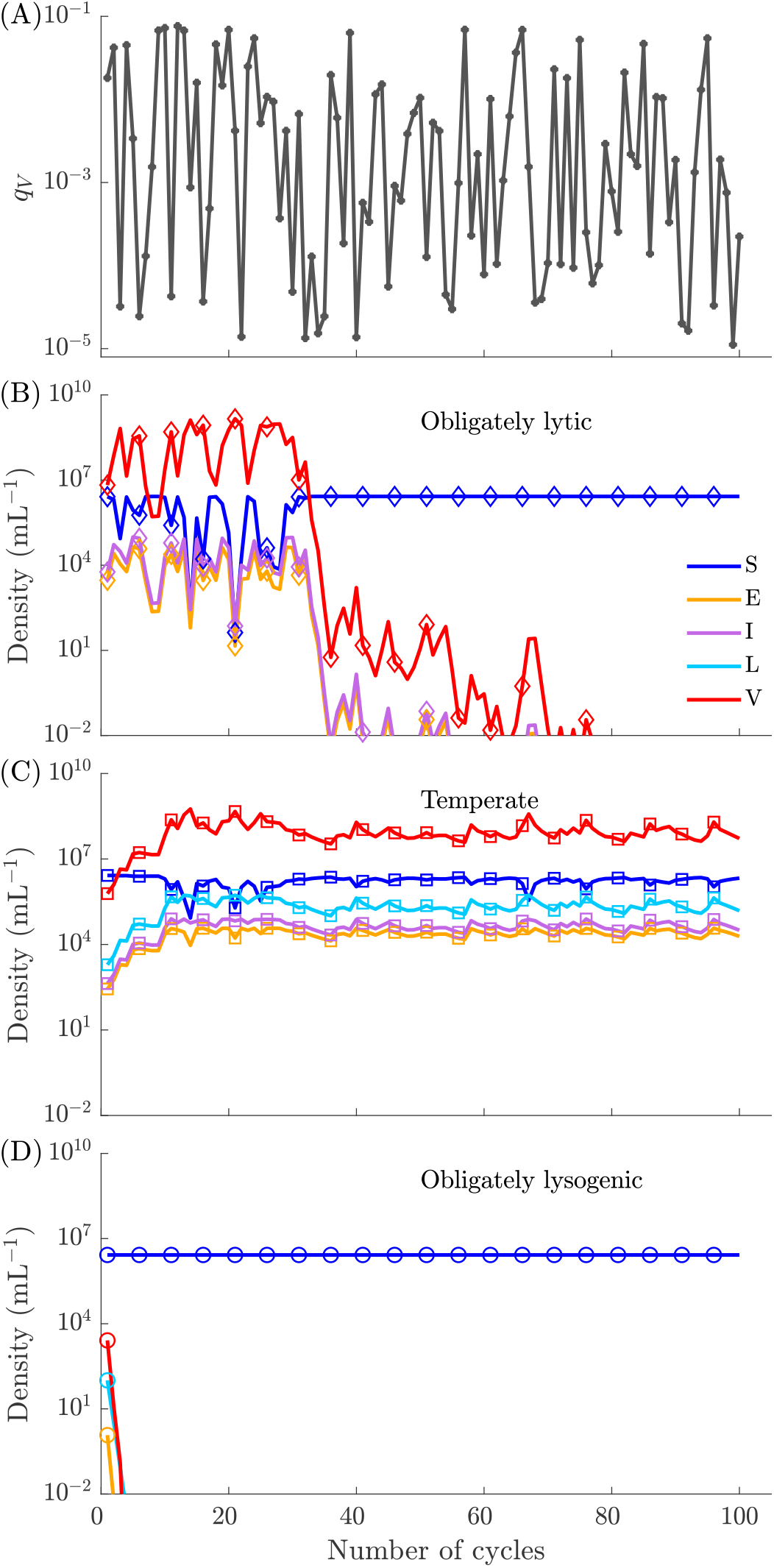
Population dynamics with stochastic filtration steps: (A) *q*_*V*_ values for a hundred growth cycles, chosen from a log uniform distribution over the span [10^−5^, 10^−1^]. Population densities at the end of each growth cycle for (B) obligately lytic (*p* = 0, *γ* = 0 hr^−1^), (C) temperate (*p* = 0.5, *γ* = 0.083 hr^−1^) and (D) obligately lysogenic (*p* = 1, *γ* = 0 hr^−1^) viruses across hundred 24 hr cycles. 10% lysogens are passaged from one cycle to the next in every case, but the virion fraction passaged from cycle-to-cycle is set by *q*_*V*_ shown in (A). Markers at every 5^th^ cycle denote that the lines represent discrete cycle-to-cycle dynamics and not continuous time dynamics. All other simulation parameters can be found in Table S1 and Table S2.

## 4 Discussion

In this study, we showed how long-term reproduction pressures (set by periodic differential dilution during filtration) acting in conflict with short-term reproduction pressures (set by host and nutrient availability) favor the emergence of temperate strategies that include both lysis and lysogeny. In particular, we showed that even if an obligately lytic virus produces more offspring over a short timescale, a temperate virus might do better on a longer timescale due to higher survival rate of lysogens. We showed that across a continuum of traits between lysis and latency, strategies that balance exploitation of hosts in the short-term and maintenance of lysogen populations in the long-term persist while obligately lytic and obligately lysogenic strategies go locally extinct. Changing the environmental context changes the eco-evolutionary outcomes of the temperate phage-host system. More latent strategies are favored when the growth cycle periods are shorter and when lysogens are passaged across growth cycles. However, lytic strategies are favored when the growth cycle periods are longer and when virions are passaged across growth cycles. Finally, temperate strategies provide resilience against environmental stochasticity. In scenarios with fixed lysogen mortality but variable virion mortality, obligately lysogenic and obligately lytic strategies are more susceptible to population crashes than temperate strategies.

The present findings extend previous work which used a cell-centric measure of fitness to identify conditions that favored latency, both in host-only conditions and in scenarios with an endemic lytic phage [28, 41–43]. These studies showed that low host availability and high phage mortality favors latency. However, the cell-centric reproduction number provides information about the short-term success of a viral strategy at the scale of an individual infected cell. In contrast, [27] used analysis of steady states to identify conditions that allowed for long-term persistence of temperate phage, showing that strong negative feedback between phage and host densities favored latency while mechanisms that decoupled host density from phage density favored lysis. Our framework allows for evaluating the realistic scenario (extending the work of [35]) that external drivers (periodic host input and differential dilution) can lead to conflicting selection pressures across timescales.

These findings came with several important caveats. For example, the present model does not incorporate phenotypic plasticity in the lysis-lysogeny switch. Across phage-bacteria systems, the probability of lysogen formation as well as the rate of prophage induction can depend on the physiological state of the host. As nutrients become scarce, host metabolism slows down, which in turn slows the infection cycle; the latent period of infection increases and burst size decreases as nutrient availability reduces [43–47]. The lysis-lysogeny switch is also sensitive to virus-host ratios, which can be sensed either through secreted molecules such as arbitrium [48, 49] or through cellular multiplicity of infection [2, 3]. Since the nutrient, host and viral densities are tightly coupled during growth cycles, phenotypic plasticity may play an important role in modulating population dynamics and evolution of viral traits [43, 47, 50, 51]. Our results suggest that temperate strategies are beneficial, even in the absence of any plasticity in response to internal/external cues regarding host or resource availability. Extending the present eco-evolutionary model to incorporate phenotypic plasticity and/or signalling cues would allow a detailed exploration of conflicts of selection across time-scales, as well as a means to evaluate the evolutionary benefits of plastic temperate strategies. In doing so, the use of a serial passage setup could make such studies experimentally tractable.

In closing, leveraging a serial passage model of the eco-evolutionary dynamics of temperate phage, we have shown how short-term selection pressures to reproduce during boom periods and long-term selection pressures to persist in bust periods can work in opposition, leading to the emergence of evolutionarily stable temperate strategies involving a mix of both lysis and lysogeny. In doing so, we also identified how ecological drivers, including the duration of growth cycles and the severity of bust periods, can tilt the balance towards strategies with more lysogenic-like traits or more lytic-like traits. The emergence of intermediate temperate phenotypes and the complexity of feedback during growth periods also points to directions that are likely amenable to further theoretical and experimental study: exploring how plasticity in viral strategies modulates ecological dynamics in the near-term and how this plasticity evolves in the long-term.

## Acknowledgements

We thank J. Harris, J. Meyer, S. Gandon and S. Lion for their inputs in model development. We also thank R. Dey for reviewing the code and suggesting improvements.

## Data availability

There is no experimental data associated with this manuscript. All code associated with the manuscript was written in MATLAB 2022b and MATLAB 2023b and is available at https://github.com/tapangoel1994/EcoEvoDynamicsInPeriodicEnvironments and is archived on Zenodo [37].

## Supplementary Material

**Figure S1.**
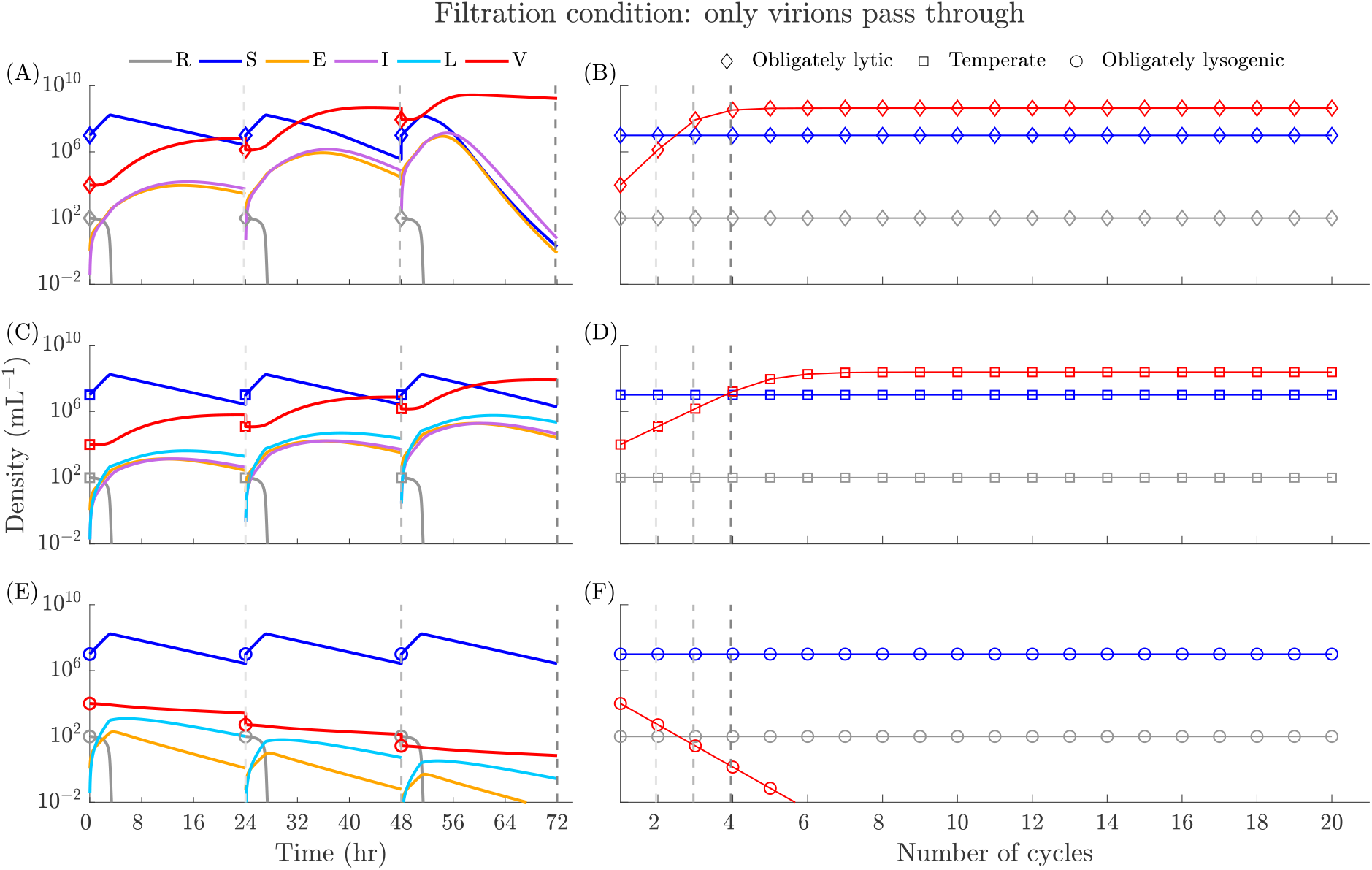
Population dynamics when only virions pass through the filter: Population dynamics over the first three 24 hr growth cycle for (A) obligately lytic (*p* = 0, *γ* = 0 hr^−1^) (⋄), (C) temperate (*p* = 0.5, *γ* = 0.083 hr^−1^) (□) and (E) obligately lysogenic (*p* = 1, *γ* = 0 hr^−1^) (◦) viruses when only virions pass through the filter (*q*_*V*_ = 0.2). Population densities at the beginning of each growth cycle for (B) obligately lytic (*p* = 0, *γ* = 0 hr^−1^) (⋄), (D) temperate (*p* = 0.5, *γ* = 0.083 hr^−1^) (□) and (F) lysogenic (*p* = 1, *γ* = 0 hr^−1^) (◦) viruses when only virions pass through the filter. The dashed vertical lines at the 24 hr, 48 hr and 72 hr marks in plots A, C and E correspond to the dashed vertical lines at cycle number 2,3 and 4 in plots B, D and F respectively.

**Figure S2.**
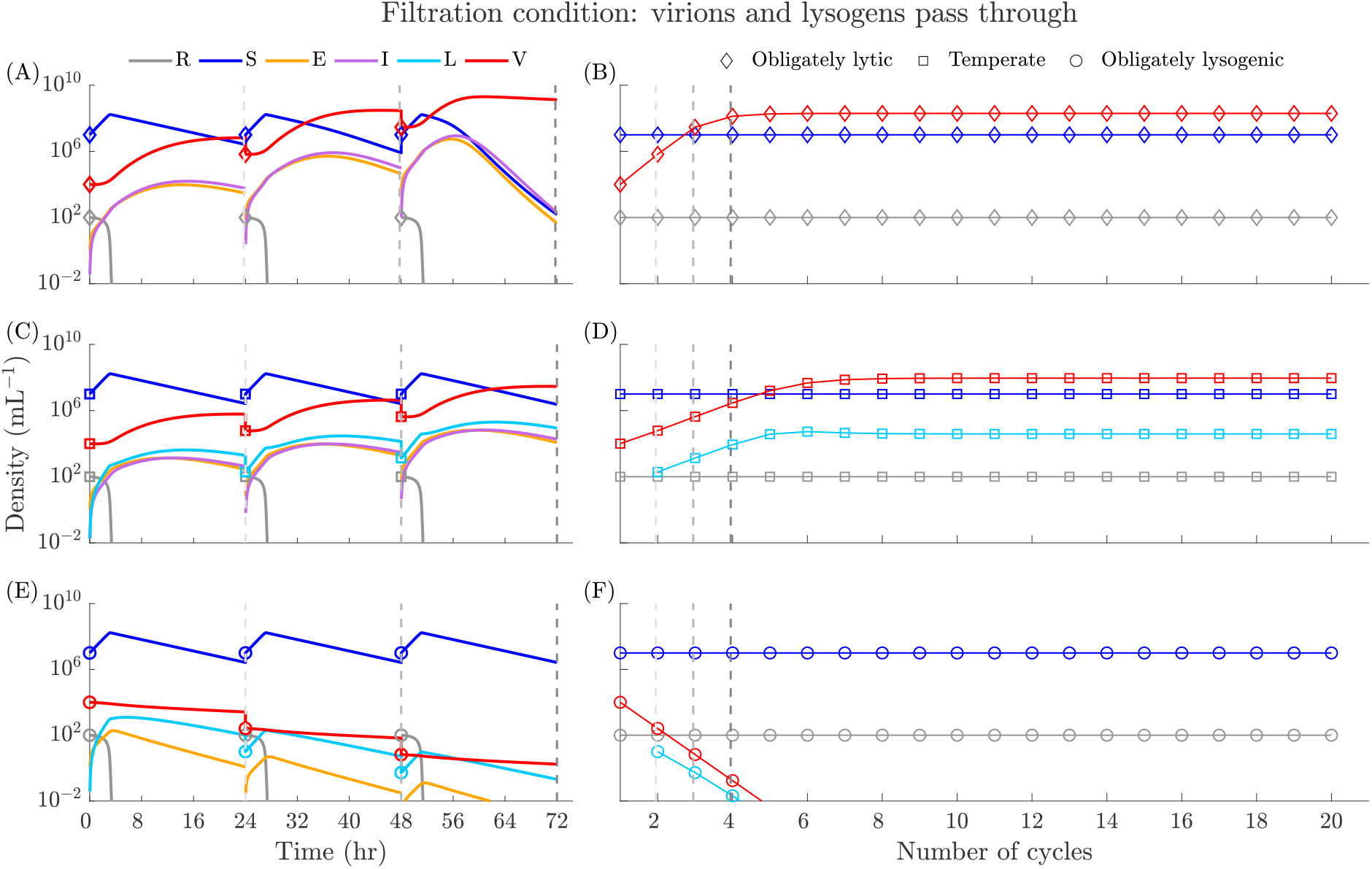
Population dynamics when virions and lysogens pass through the filter: Population dynamics over the first three 24 hr growth cycle for (A) obligately lytic (*p* = 0, *γ* = 0 hr^−1^) (⋄), (C) temperate (*p* = 0.5, *γ* = 0.083 hr^−1^) (□) and (E) obligately lysogenic (*p* = 1, *γ* = 0 hr^−1^) (◦) viruses when virions and lysogens pass through the filter (*q*_*L*_ = 0.1, *q*_*V*_ = 0.1). Population densities at the beginning of each growth cycle for (B) obligately lytic (*p* = 0, *γ* = 0 hr^−1^) (⋄), (D) temperate (*p* = 0.5, *γ* = 0.083 hr^−1^) (□) and (F) lysogenic (*p* = 1, *γ* = 0 hr^−1^) (◦) viruses when virions and lysogens pass through the filter. The dashed vertical lines at the 24 hr, 48 hr and 72 hr marks in plots A, C and E correspond to the dashed vertical lines at cycle number 2,3 and 4 in plots B, D and F respectively.

**Figure S3.**
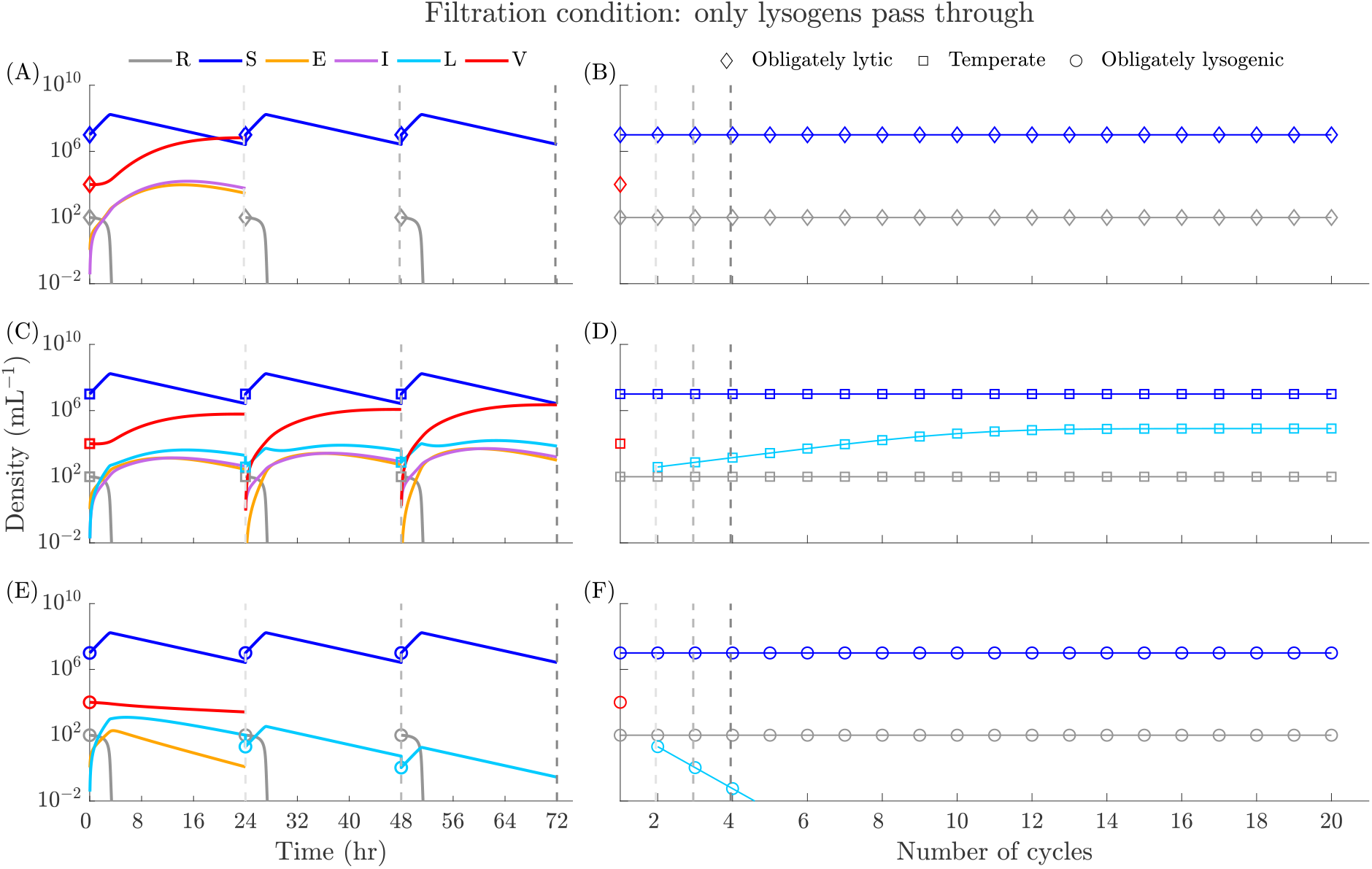
Population dynamics when only lysogens pass through the filter: Population dynamics over the first three 24 hr growth cycle for (A) obligately lytic (*p* = 0, *γ* = 0 hr^−1^) (⋄), (C) temperate (*p* = 0.5, *γ* = 0.083 hr^−1^) (□) and (E) obligately lysogenic (*γ* = 0 hr^−1^, *p* = 1) (◦) viruses when only lysogens pass through the filter (*q*_*L*_ = 0.2). Population densities at the beginning of each growth cycle for (B) obligately lytic (*p* = 0, *γ* = 0 hr^−1^) (⋄), (D) temperate (*p* = 0.5, *γ* = 0.083 hr^−1^) (□) and (F) lysogenic (*p* = 1, *γ* = 0 hr^−1^) (◦) viruses when only lysogens pass through the filter. The dashed vertical lines at the 24 hr, 48 hr and 72 hr marks in plots A, C and E correspond to the dashed vertical lines at cycle number 2, 3 and 4 in plots B, D and F respectively.

**Figure S4.**
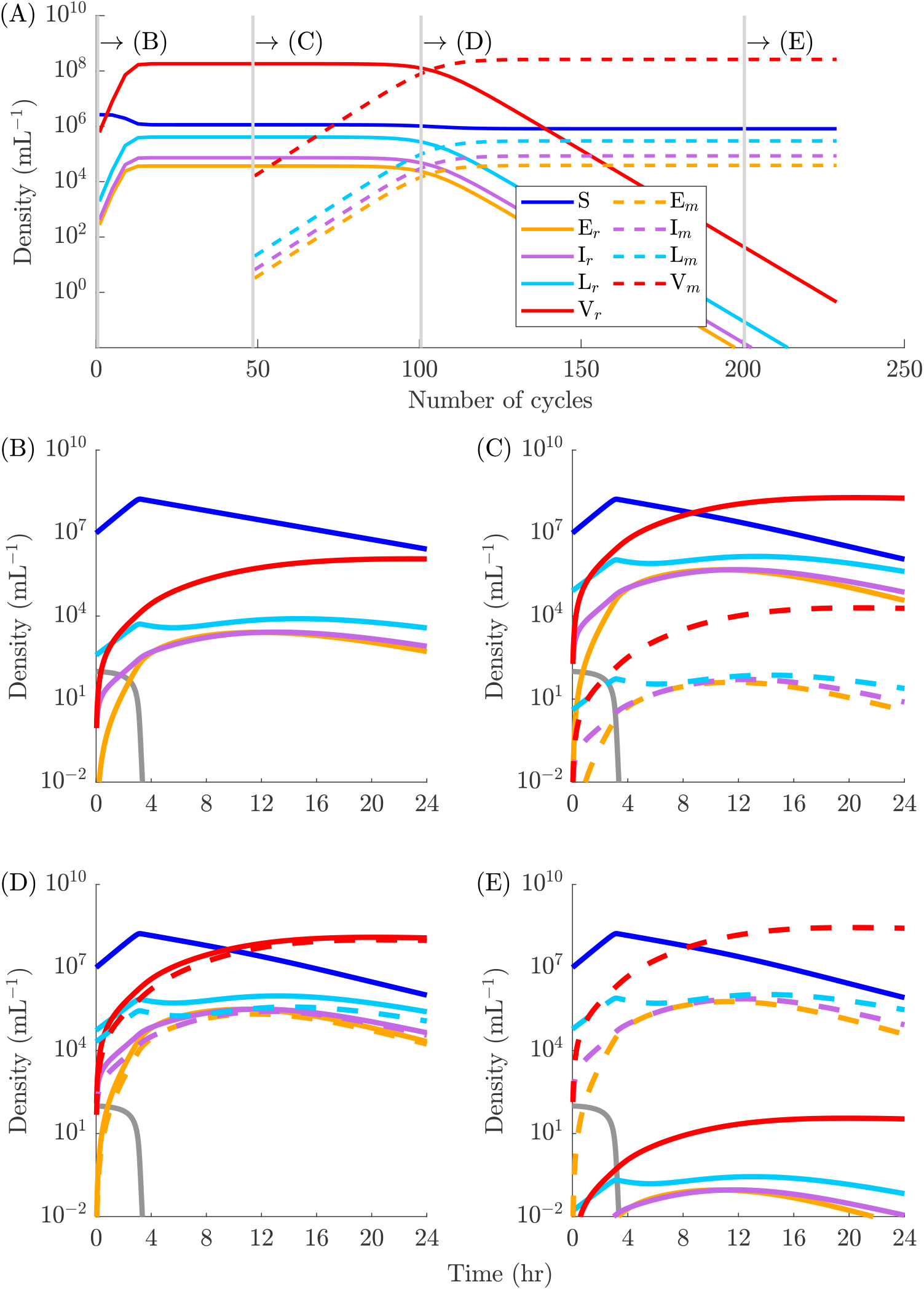
Cycle-to-cycle invasion dynamics: (A) Final cell/virion densities at the end of each cycle for (r)esident (*p*^*^ = 0.52, *γ*^*^ = 0.083 hr^−1^) and (m)utant (*p*_ESS_ = 0.3, *γ* = 0.083 hr^−1^) virus types. The growth cycle duration is 24 hr, and only 20% of lysogens (*q*_*L*_ = 0.2) from one growth cycle pass to the next growth cycle. Gray vertical bars highlight the 1st, 49th, 100th and 200th growth cycles, the within cycle dynamics for which are shown in panels (B), (C), (D) and (E) respectively.

**Figure S5.**
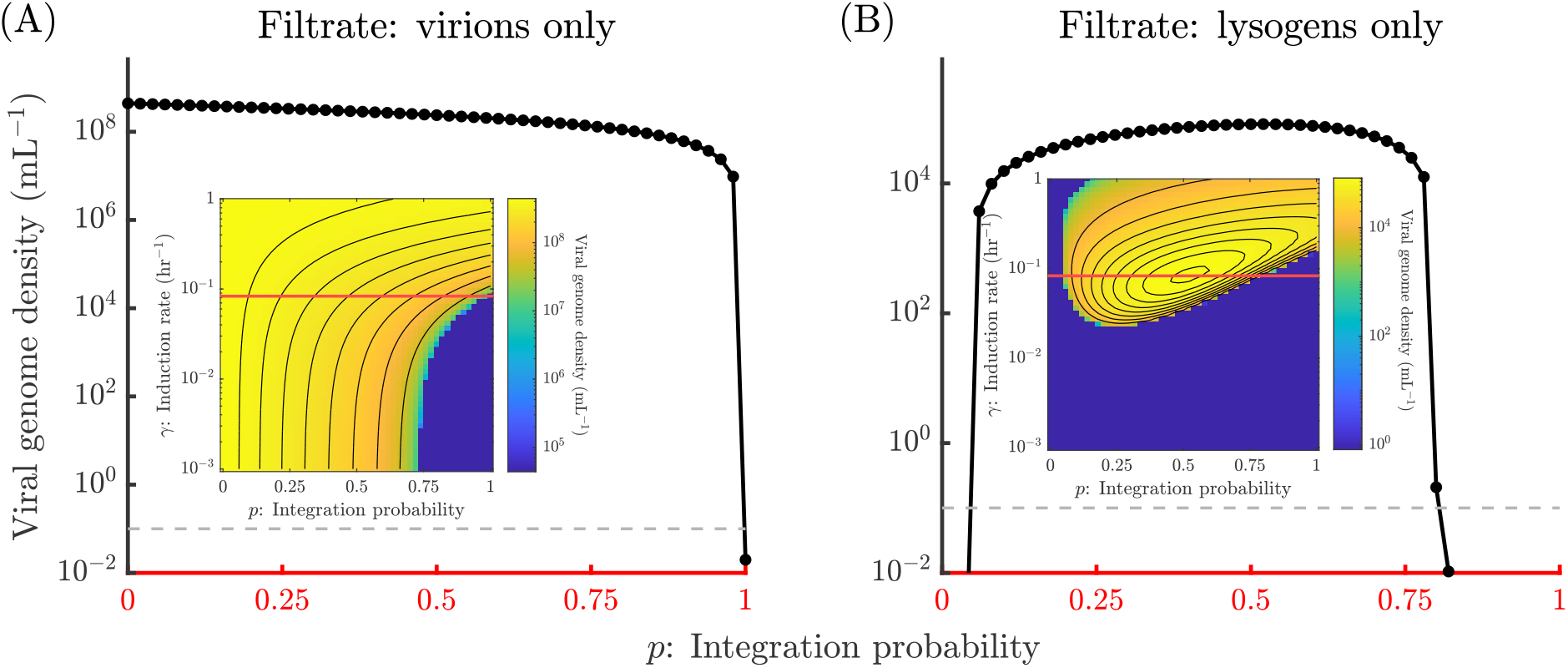
Steady state viral genome densities for a fixed induction rate: Density of viral genomes at the beginning of a steady state growth cycle as a function of integration probability for a fixed induction rate (*γ* = 0.083 hr^−1^) when (A) only virions (*q*_*V*_ = 0.2) and, (B) only lysogens (*q*_*L*_ = 0.2) pass through the filter. The cycle duration is *T* = 24 hr. The insets are the steady state viral genome density heatmaps in Figure 4. The horizontal red lines in the insets correspond to the x-axes of (A) and (B). Steady state viral density is assumed to be too small for invasion analysis if it is below the dashed gray line *y* = 100*ϵ*.

**Figure S6.**
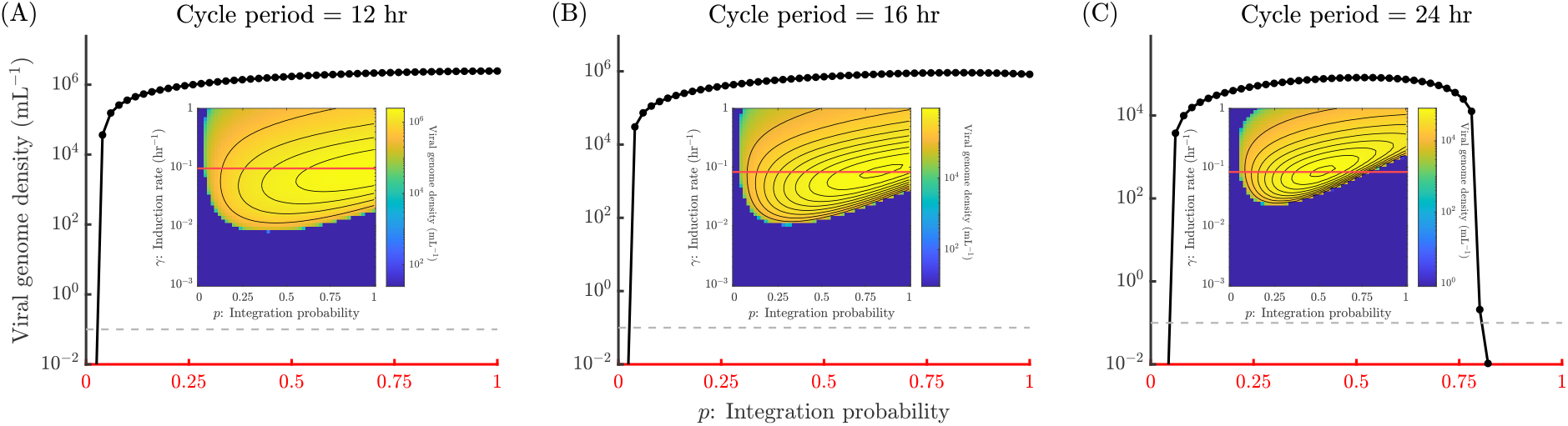
Steady state viral genome densities for a fixed induction rate: Density of viral genomes at the beginning of a steady state growth cycle as a function of integration probability for a fixed induction rate (A) *γ* = 0.095 hr^−1^, (B) *γ* = 0.083 hr^−1^ and (C) *γ* = 0.083 hr^−1^ when only lysogens (*q*_*L*_ = 0.2) pass through the filter. The growth cycle durations are *T* = 12 hr in (A), *T* = 16 hr in (B) and, *T* = 24 hr in (C). The insets are the steady state viral genome density heatmaps in Figure 5. The horizontal red lines in the insets correspond to the x-axes of (A), (B) and (C). Steady state viral density is assumed to be too small for invasion analysis if it is below the dashed gray line *y* = 100*ϵ*.

**Figure S7.**
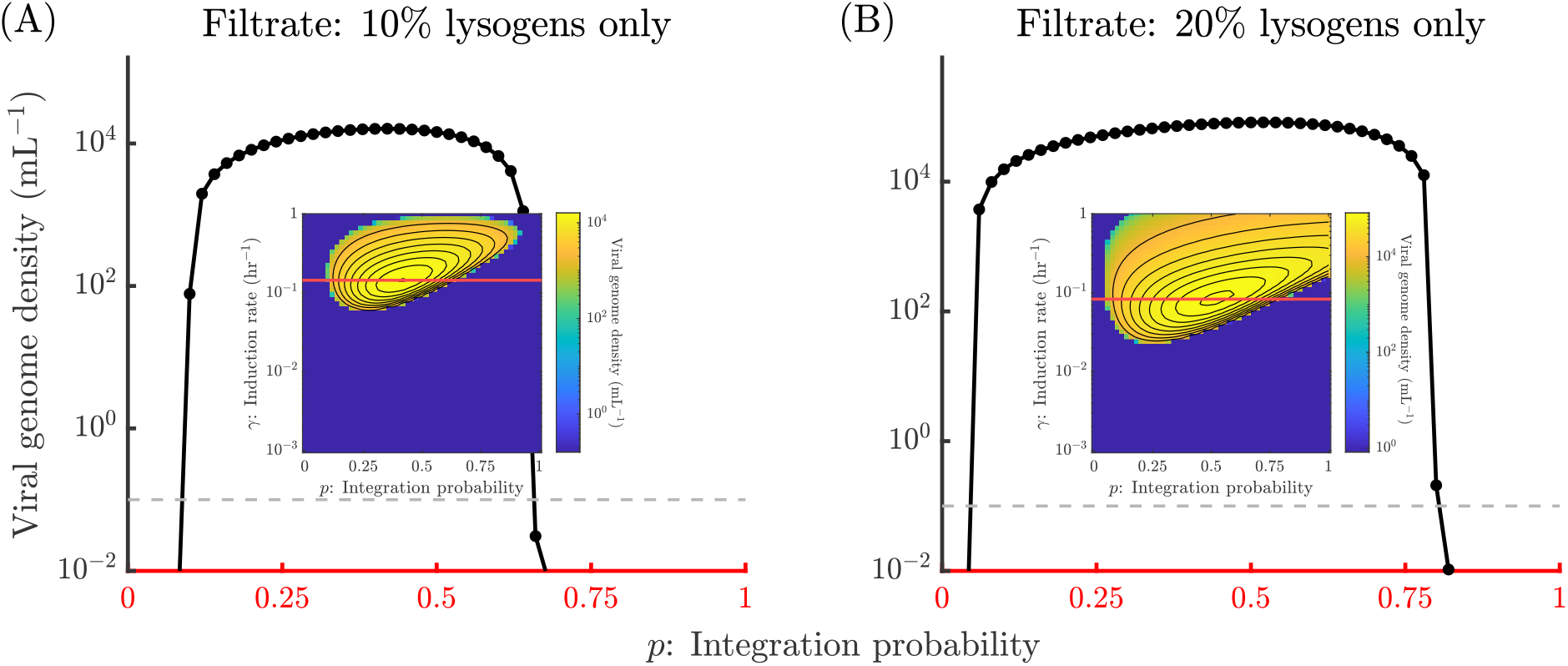
Steady state viral genome densities for a fixed induction rate: Density of viral genomes at the beginning of a steady state growth cycle as a function of integration probability for a fixed induction rate (A) *γ* = 0.144 hr^−1^ when 10% lysogens (*q*_*L*_ = 0.1) pass through the filter, and (B) *γ* = 0.083 hr^−1^ when 20% lysogens (*q*_*L*_ = 0.2) pass through the filter. The cycle duration is *T* = 24 hr for both (A) and (B). The insets are the steady state viral genome density heatmaps in Figure 6. The horizontal red lines in the insets correspond to the x-axes of (A) and (B). Steady state viral density is assumed to be too small for invasion analysis if it is below the dashed gray line *y* = 100*ϵ*.

## Tables

**Table S1.** SEILV model parameters (see [49], Table A1).

| Parameter name | Parameter symbol | Value | Units |
| --- | --- | --- | --- |
| Conversion efficiency | $e$ | $5 \times 10^{-7}$ | $\mu\text{g}/\text{cell}$ |
| Maximal cell growth rate | $\mu_{max}$ | 1.2 | $\text{hr}^{-1}$ |
| Half velocity constant | $R_{in}$ | 4.0 | $\mu\text{g}/\text{mL}$ |
| Death rate of susceptible host | $d_S$ | 0.2 | $\text{hr}^{-1}$ |
| Death rate of exposed cells | $d_E$ | 0.2 | $\text{hr}^{-1}$ |
| Death rate of lysogens | $d_L$ | 0.2 | $\text{hr}^{-1}$ |
| Death rate of lytically infected cells | $d_I$ | 0.2 | $\text{hr}^{-1}$ |
| Transition rate | $\lambda$ | 2 | $\text{hr}^{-1}$ |
| Lysis rate | $\eta$ | 1 | $\text{hr}^{-1}$ |
| Burst size | $\beta$ | 50 | |
| Adsorption rate | $\phi$ | $3.4 \times 10^{-10}$ | $\text{mL}/\text{hr}$ |
| Viral decay rate | $m$ | 1/24 | $\text{hr}^{-1}$ |
| Integration probability | $p$ | $\{0, 0.02, 0.04 \dots 0.98, 1\}$ | |
| Induction rate | $\gamma$ | $\{10^{-3}, \dots 10^{-0.06}, 10^0\}$ | $\text{hr}^{-1}$ |

**Table S2.** Initial conditions and filtration conditions for serial passage simulations.

| Parameter name | Parameter symbol | Value | Units |
| --- | --- | --- | --- |
| Initial resource density | $R_0$ | 100 | $\mu\text{g}/\text{mL}$ |
| Initial host density | $S_0$ | $10^7$ | $\text{cells}/\text{mL}$ |
| Initial virion density | $V_0$ | $10^4$ | $\text{virions}/\text{mL}$ |
| Initial virion density (resident) | $V_{a0}$ | $10^4$ | $\text{virions}/\text{mL}$ |
| Initial viral genome density (mutant) | $M$ | $10\epsilon$ | $\text{genomes}/\text{mL}$ |
| Fraction of resources in filtrate | $q_R$ | 0 | |
| Fraction of susceptible hosts in filtrate | $q_S$ | 0 | |
| Fraction of exposed cells in filtrate | $q_E$ | 0 | |
| Fraction of lytically infected cells in filtrate | $q_I$ | 0 | |
| Fraction of lysogens in filtrate | $q_L$ | $\{0, 0.1, 0.2\}$ | |
| Fraction of virions in filtrate | $q_V$ | $\{0, 0.1, 0.2\}$ | |
| Duration of growth cycle | $T$ | $\{12, 16, 24\}$ | hr |
| Critical density threshold | $\epsilon$ | $1 \times 10^{-3}$ | $\text{mL}^{-1}$ |

